# A simple and dynamic thermal gradient device for measuring thermal performance in small ectotherms

**DOI:** 10.1101/2020.08.28.271767

**Authors:** Marshall W. Ritchie, Jeff W. Dawson, Heath A. MacMillan

## Abstract

The body temperature of ectothermic animals is heavily dependent on environmental temperature, impacting fitness. Laboratory exposure to favorable and unfavorable temperatures is used to understand these effects, as well as the physiological, biochemical, and molecular underpinnings of variation in thermal performance. Although small ectotherms, like insects, can often be easily reared in large numbers, it can be challenging and expensive to simultaneously create and manipulate several thermal environments in a laboratory setting. Here, we describe the creation and use of a thermal gradient device that can produce a wide range of constant or varying temperatures concurrently. This device is composed of a solid aluminum plate and copper piping, combined with a pair of programmable refrigerated circulators. As a simple proof-of-concept, we completed single experimental runs to produce a low-temperature survival curve for flies (*Drosophila melanogaster*) and explore the effects of daily thermal cycles of varying amplitude on growth rates of crickets (*Gryllodes sigillatus*). This approach avoids the use of multiple heating/cooling water or glycol baths or incubators for large-scale assessments of organismal thermal performance. It makes static or dynamic thermal experiments (e.g., creating a thermal performance or survival curves, quantifying responses to fluctuating thermal environments, or monitoring animal behaviour across a range of temperatures) easier, faster, and less costly.

## Introduction

The thermal environment can directly impact organismal survival and fitness, and small ectotherms can respond to changing thermal environments by altering behaviour or physiology within the lifetime of an individual or over evolutionary time [1,2]. Our understanding of how animals respond to their thermal environment has come largely from studies focused on small ectothermic animals with short generations times that can be reared rapidly and inexpensively in the laboratory. As insects are easily manipulated and maintained in the lab, model insect species are widely used for studies of the genetics, molecular biology, environmental physiology, behaviour, and evolution of thermal performance traits [3–7]. Laboratory studies of *Drosophila* thermal tolerance, for example, have recently allowed for investigation of the molecular or physiological processes that limit their biogeography, and how limits to thermal performance may evolve following changes in abiotic conditions [4,8–10].

A common measure of thermal tolerance is survival following high or low-temperature exposure. Survival experiments are typically assessed using one of two different experimental designs. The first is prolonged exposure to a single lethal temperature [11,12]. This approach is limited to the selected exposure temperature, which may or may not be the ideal temperature to measure the thermal tolerance of the chosen species, or discriminate variation in thermal tolerance among treatment groups. An alternative approach is to use a range of exposure temperatures [13–16]. This approach is used to generate thermal tolerance or performance curves by examining, for example, the effects of temperature on survival [17], growth rates [18], reproductive capacity [8], motor performance [19] or behaviour [20]. In these experiments, different individuals are exposed to different static temperatures to examine underlying temperature effects on life history and to understand better how prior thermal experiences might influence these relationships [19,21,22].

In nature, animals do not experience a drastic increase or decrease in temperature but rather experience gradual temperature increases or decreases as well as non-linear and stochastic temperature profiles [2,23]. These predictable or unpredictable changes in temperature affect insects differently than would a sudden increase or decrease in temperature applied in a typical lab setting [23]. Experiments involving ramping temperatures are now common [24–26], and there is concern over the effects of the rate of temperature change on the experimental outcome. In light of this concern, some authors have directly measured the effects of different ramping rates on thermal performance traits [27–29], choosing a handful of ramping rates to test. Ramping temperatures are just one example that demonstrates a growing appreciation for thermal performance and evolution studies that embrace complexity of natural thermal environments as well as the complexity of physiological responses to those environments [30]. Experimental design, however, can strongly influence research outcomes, and several factors limit our ability to test a wide range of conditions at once.

When performing an experiment that requires multiple constant temperatures or programmed temperature changes over time, it can be challenging to maximize the number of different treatments available because of limited equipment and keep unintended sources of variation among treatments to a minimum. Variation in the resulting data from such experiments can come from either using multiple heating/cooling devices all running at different temperatures or having to use individual specimens from multiple successive generations or of varying age because of limited capacity or versatility of the equipment or time. While accounting for these variables statistically is sometimes an option, controlling for them is preferred. In addition to potentially impacting data quality, equipment limitations can significantly increase the resources and time required to perform a thermal experiment, thereby restricting both data quality and quantity.

Here, we describe the manufacturing and testing of a thermal gradient plate and associated thermal bath setup that can streamline thermal experiments. We believe that our design allows for the rapid and accurate quantification of a variety of organismal thermal tolerance or performance metrics. By connecting two heated and/or refrigerated circulating baths to either end of a custom aluminum plate, stable and predictable thermal gradients can be formed (documented here and previously with various designs, see: [31,32]). If one or both of those circulating baths can also be programmed, however, a single setup can become a powerful tool for examining the impacts of both static and dynamic thermal conditions on organismal performance and fitness. We describe the utility of this system from our perspective as insect thermal biologists but small organisms or samples such as other terrestrial or aquatic invertebrates, plants, unicellular eukaryotes, bacteria, cell cultures, or even enzymes could be studied with this approach. Labor-intensive experiments can be easily and rapidly accomplished using the described system in a single run. As a proof-of-concept, we generated low-temperature survival curves of male and female *D. melanogaster* (typically produced using multiple cooling baths) and examined how the amplitude of diurnal temperature cycles influenced the growth rates of tropical house crickets (*Gryllodes sigillatus*).

## Materials and Methods

### Building the plate

The thermal plate was created using parts (Table 1) that were assembled in the laboratory. These pieces are readily available from a hardware store and/or publicly accessible metal supplier or machine shop. The gradient device was built using a 91.4 cm (36”) x 45.7 cm (18”) x 2.5 cm (1”) solid aluminum plate (Figure 1A). We milled channels on each side and at both ends of the plate (12 channels in total; Figure 1B). Given the scale of the plate, this milling was the most technically demanding part of the build but could be easily completed by a local machine shop if necessary. Each channel had dimensions of 0.95 cm (3/8”) x 0.95 cm (3/8”) and sets of 3 channels were milled 0.95 cm (3/8”) apart. Copper tubing (0.95 cm thick (3/8” OD) x 49 cm (c. 19.25”) long) was gently hammered into the channels, being careful not to damage the tubing in any way that would block fluid flow or cause a leak. Hammering was done until the copper tubing was flat with the surface of the plate (Figure 1A). We left 1.65 cm (c. 0.5”) of tubing overhang on each side of the plate for plumbing connections (Figure 1C).

**Table 1:**
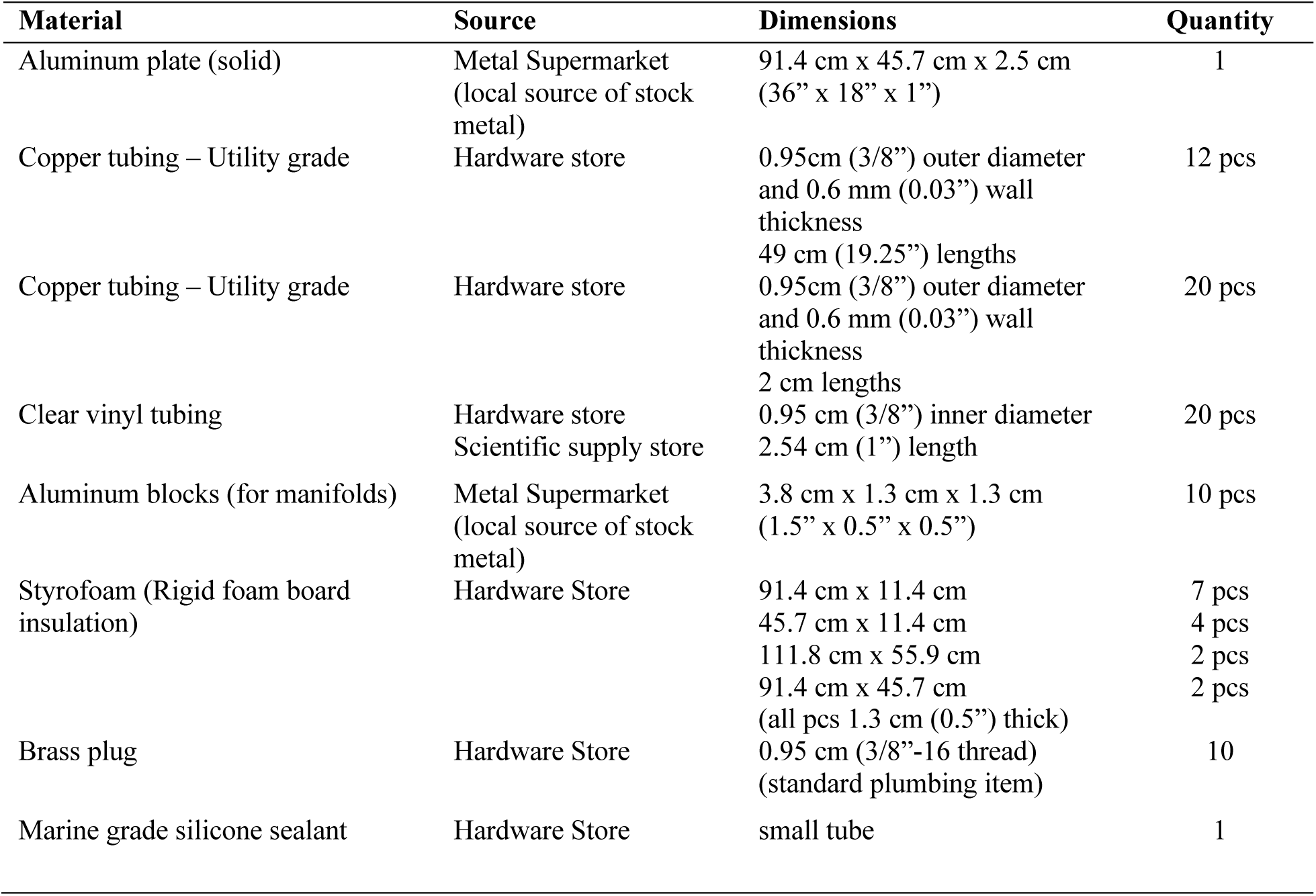
List of material to assemble the plate with the dimensions of each item as well as the number required. In addition to the other materials, two refrigerated circulating baths were used. These can be programmable or non-programmable depending on the experimental design. All programs shown herein were conducted with one programmable and one non-programmable bath (capable of static temperatures only).

**Figure 1:**
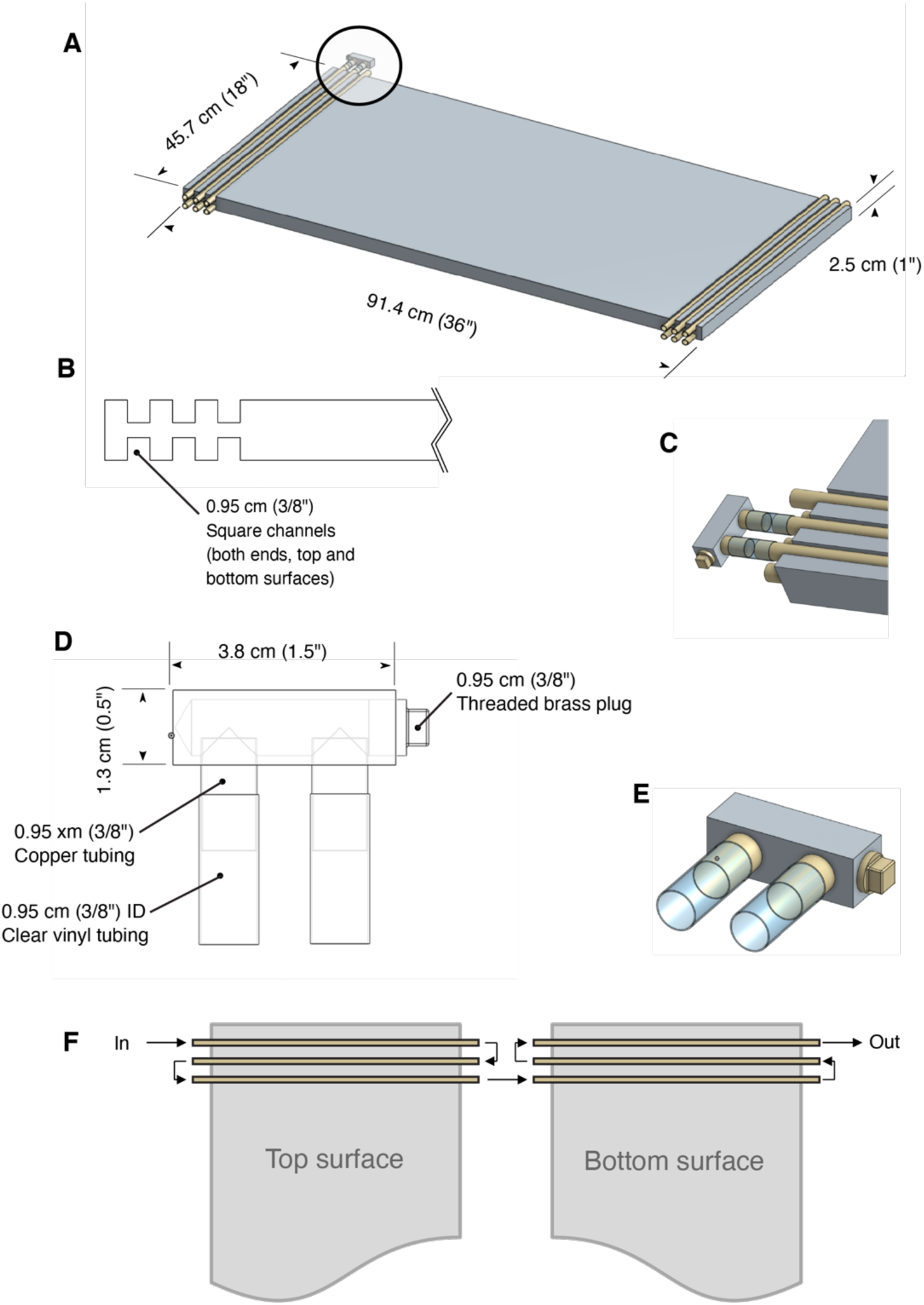
Design and dimensions of the thermal gradient plate. A) Assembled plate and copper tubing with example manifold shown in place (circled region). B) Side view of milled square channels in aluminum plate. C) Manifold connection to exposed copper pipe ends using clear vinyl tubing. D) Manifold design from aluminum block, copper pipe, brass plug, and tubing. E) Close-up of example manifold. F) Direction of fluid flow through the system once all manifolds are connected (manifolds not shown for clarity).

After several attempts, we could not bend our copper tubing into a ‘U’ shaped configuration without damaging it and therefore opted to construct simple manifolds to complete the plumbing circuit (Figure 1D). Aluminum blocks (3.8 cm (1.5”) x 1.3 cm (1.5”) x 1.3 cm (1/2”)) were used to create the custom manifolds. Two 0.95 cm (3/8”) holes were drilled adjacent to each other in the long face of the block (Figure 1D). The spacing of the holes aligned with the spacing of the adjacent channels in the aluminum block. Another hole was drilled in one end face deep enough into the block so as to create a cavity connecting the first two holes (Figure 1D). This hole was drilled with a 5/16” drill bit to allow the hole to be tapped to accommodate a 3/8”-16 threaded brass plug. Two shorter lengths (c. 2 cm) of 0.95 cm (3/8” OD) copper tubing were press-fit into the adjacent holes and a 3/8”-16 brass plug was installed in the threaded end-face hole using sealant to prevent leaking when fluid was under pressure. The manifolds were then connected to the ends of the copper tubing in the aluminum plate with 2.5 cm (c. 1”) lengths of 0.95 cm (3/8” ID) clear vinyl tubing (Figure 1C). We additionally secured the vinyl tubing connections with steel wire.

A benefit of using the manifolds attached to the main plate via short lengths of tubing was to compensate for slight variations in spacing of the channels and tubing, and to avoid the possibility of the seals breaking with thermal expansion of the plate.

The remaining two ends of the copper tubing at each end of the plate were attached to their respective cooling baths. By connecting the manifolds to the plate in a specific arrangement fluid was directed through all 6 of the lengths of tubing at each end of the plate (Figure 1F). Extruded polystyrene foam board (rigid foam-board insulation) was then cut to fit all sides of the plate to leave an approximately 7.5 cm (3”) airspace between the top of the aluminum plate and the bottom of the lid. The air gap allowed space for the placement of samples or metal tins containing samples (described below).

### System testing

The plate was tested to ensure that a stable and reproducible temperature gradient could be formed on the surface of the aluminum plate. First, eight type-K thermocouples were equally spaced along the length of the plate at the mid-line of the plate width (Figure 2A). The positioning of each probe was pseudo-randomized during each replicate to minimize bias from using the same probe in the same location. Three different thermal gradients were independently tested to examine the functional range of the device. Three gradients 0°C to +10°C, 0°C to +25°C, and 0°C to +40°C) were each tested three times, for 2 h (each replicate). We also measured temperatures across the width of the plate at the same locations across the length of the plate.

**Figure 2:**
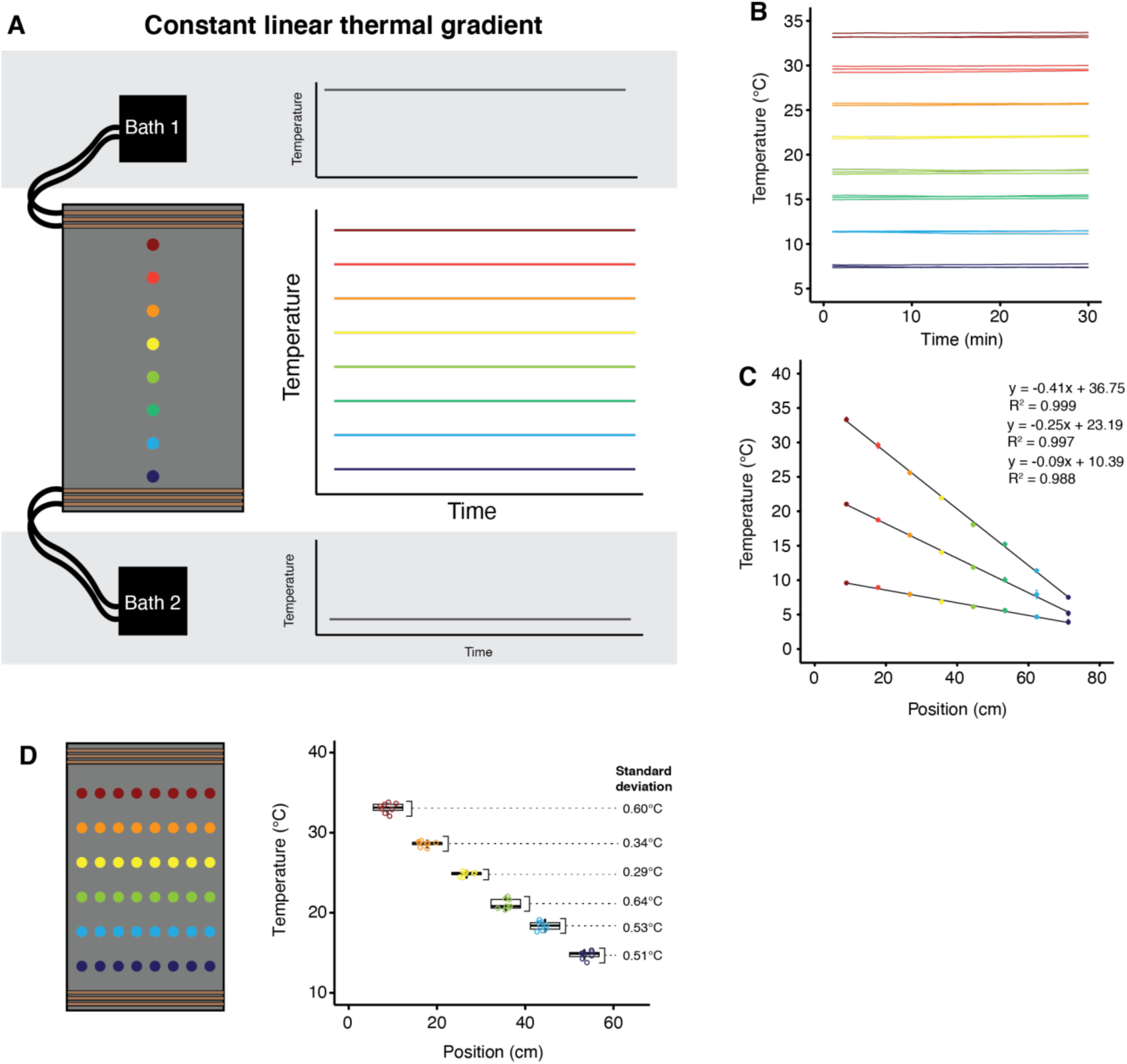
Production of stable linear gradients along the length, but not the width of the gradient plate. A) Probe positions and expected outcomes from establishing constant temperatures at either end of the gradient plate. B) Actual data recorded at eight locations in the position of the plate with an established linear gradient. Circulating baths connected to the plate were set to +40°C (bath 1) and 0°C (bath 2). Each line represents a single type-K thermocouple recording over a 30 min period (n=3 per position). C) Temperatures recorded at eight locations along the length of the plate while maintaining a stable gradient. Solid circles represent the mean of three replicant measurements at each location (open circles, barely visible because of low variance). Note that the R^2^ of the temperature-position relationship is close to 1 (0.988 to 0.999). D) Measurements of temperature along the width of the plate at several points in the length of the plate with a thermal gradient of 0 to +40°C. Colors denote position along the length of the plate, and eight temperature recordings across the width of the plate were made for each position along its length. Boxplots show the variation of temperature across the plate width.

We tested the plate using three different dynamic programs that were selected based on their utility to insect thermal biologists. The first was two ramping tests from +20°C to +40°C over 1 h and 2 h at one end of the plate while the other was held constant at +20°C (simulating a ramp to acute heat stress). The second dynamic test was five cycles of +10°C to +30°C at one end while the other end was held constant at +20°C. To simulate rapid thermal cycles, this cycling was left to repeat five times, and each cycle lasted a total of two hours from +10 to +30°C and vice versa, but diurnal cycles as used for our cricket experiment (below) are commonly longer. The third dynamic test was a fluctuating thermal regime (FTR). The FTR was done by forming a 0 to +25°C gradient during a cool period, followed by an identical warm period across the plate at +25°C. FTR programs are commonly used and studied in the context of long-term insect storage [33–35]. Each of these programs was tested and recorded once.

We next turned our attention to the practical use of the plate. We tested whether small animal containers (glass *Drosophila* vials), could be placed directly on the plate, or if an intermediate containment vessel was needed to achieve stable temperatures inside the containers. We expected issues to be most pronounced with temperature gradients across a sample container that deviated far from room temperature, so these tests were conducted at low temperatures. With a gradient of −10 to +7°C formed across the plate, glass vials (25 mm diameter x 95 mm height) were placed at the −10°C end of the plate. We used type-K thermocouples to measure the air in the vial, and the bottom of the glass along with the plate. Since this experiment indicated that contact with the plate alone was not sufficient to establish stable homogeneous temperatures inside sample vials, we devised an alternative approach. We opted to use narrow troughs containing a mixture of ethylene glycol and water. We created the troughs (39.37 x 7.62 x 7.62 cm) using sheet metal (0.9 mm thick), rivets, and silicone caulking. The troughs were placed on the plate with the vials which were then placed in glycol in the troughs. This approach produced stable and homogenous temperatures throughout the glass surface of the vial as well as the air inside the vial.

### Proof of concept 1: Drosophila cold tolerance

The population of *Drosophila melanogaster* used [36] were reared in 200 mL plastic bottles containing 50 mL of a banana-based diet (containing primarily banana, active yeast, corn syrup, and barley malt). Flies were kept in an incubator at +25°C in a 12 h:12 h light/dark cycle. Flies were allowed to lay fresh eggs by transferring ∼800 adult flies into a population cage containing food in a petri dish and were left for 24 h before being removed. This process resulted in roughly 1200 eggs laid in each cage. The food containing the eggs was divided up and placed into fresh glass vials (approx. 100 eggs per vial).

All flies were sorted by sex on the day of adult emergence under light CO_2_. Flies were not exposed to CO_2_ for more than 10 min to ensure no long-term physiological effects [37,38]. Females were transferred in groups of 10 into glass vials with 7 mL of fresh banana food (mainly made up of bananas, corn syrup, agar, yeast).

Our sheet metal troughs (described above) were laid across the width of the plate at nine positions across the length of the plate. Each tin was filled with 1 L of a mix of ethylene glycol and water and was placed on the plate once a stable −6°C to −1°C gradient had formed. Nine metal tins were placed on the plate from one end to the other, which resulted in each vial of flies being exposed to one of nine temperature points, depending on location. Type-K thermocouples were placed at the bottom of two vials in each tin to record temperatures experienced by the flies. Adult flies were transferred to empty glass vials and restricted to the bottom 25% of the vial with a foam stopper. All of the vials were then placed into the metal tins. After 4 h in the cold, flies were removed and transferred to vials with banana food and left to recover at +25°C. At 2 h, 24 h, and 48 h (after removal from the cold), survival was checked and scored based on whether the fly was able to stand upright at room temperature. Using the survival scores from 2 h after the cold stress, we tested for an effect of temperature on the ability of flies to survive at different temperatures using a generalized linear model with temperature and sex as factors. This model had a binomial error distribution and a logit-link. We then extracted data from flies exposed to - 2.21°C and used a mixed effects model (with sex and recovery time as factors and vial as a random effect) to examine how rates of survival changed over time after removal from the cold.

### *Proof of concept 2: Gryllodes sigillatus* cycling growth

The *Gryllodes sigillatus* eggs used for the experiment came from Entomo Farms, in Norwood, ON, Canada. The eggs were laid at the farm and were transferred to a +25°C incubator at day one of egg development. Hatchlings that all emerged within a 24 h period were transferred to a plastic container (24.5 cm × 12.5 cm × 19.5 cm) where they were kept at +33°C with food, shelter and water and were maintained for two weeks. Each individual was then weighed to the nearest µg using a microbalance (Sartorius ME5 model) and transferred to a 30 mm petri dish. Each petri dish contained a 200 µL PCR tube lid filled with cricket diet, a 200 µL microcentrifuge tube with water and stoppered with cotton, and a folded piece of paper for shelter. Crickets in their respective Petri dishes were placed directly on the plate at 10 different locations across the length of the plate in replicates of 10, resulting in a total sample size of 100 crickets. The treatment groups (defined by location on the plate) were divided by Styrofoam barriers to minimize airflow across the plate surface. A light strip was attached to the inside of the Styrofoam lid and was set at a 12:12 h day and night cycle (lights on at 8 AM).

One side of the plate was held at a constant 30°C while the other side was cycled daily between 20°C and 40°C. The ramp rate was set to increase or decrease by 10°C over 6 h, and a full cycle was completed every 24 h. Temperatures experienced were measured by placing two type-K thermocouples into empty cricket containers at the far ends of each treatment group. This created 10 treatment levels of daily thermal variability experienced by the crickets with those closest to the cycling end experiencing the widest range of temperatures (+38°C to +22°C).

Water for the crickets was replaced on days 2 and 4 of the 5-day cycling period to ensure the animals were not water stressed. Once 5 cycles had been completed, the crickets were removed from the plate, and the body mass of each individual was measured. The effect of thermal cycle amplitude and initial mass on final mass and the effect of thermal cycle amplitude on proportion of mass gained over the five-day period were tested using general linear models.

## Results and Discussion

### Simple thermal gradients

Based on our design, we predicted that we could achieve linear thermal gradients across the plate that were stable (Figure 2A). Indeed, stable thermal gradients formed on the gradient plate when one refrigerated circulator was set to 0°C and the other was set to +10°C, +25°C or +40°C (Figure 2B). The relationship between distance (across the plate) and temperature always closely fit simple linear models (R^2^ between 0.988 and 0.999; Figure 2C). The intercepts of each of the gradients were close to the value of the highest set temperature (Figure 2C).

In a separate run using circulator settings of constant 0°C and 40°C, we examined the reliability of temperatures across the width of the plate. The temperature across the width of the plate was measured to ensure consistent temperatures would be experienced by an organism at the same position across the length of the plate (Figure 2D). The temperature measured across the width of the plate was consistent at each location across the length (Figure 2D). The standard deviation of temperature at each of the positions across the width of the plate varied between 0.3 and 0.6°C (Figure 2D), which is well within the expected error rate of type-K thermocouples (∼1.1°C or 0.4%). Thus, like previously described thermal gradient plates [32], our system can produce linear temperature gradients across the length of the plate, and these temperatures are stable over time (Figure 2).

### Dynamic thermal gradients

The first dynamic gradient we attempted was an upward temperature ramp with one bath while the other was held at a constant temperature (Figure 3A). This dynamic gradient was repeated twice with a 2 h (0.17°C min^-1^ rate) and 1 h (0.33°C min^-1^ rate) ramp being completed by the bath. Both of these experiments resulted in a range of ramping rates across the plate that varied linearly with position (Figure 3D, G). Thus, this system can easily be used to study the effects of temperature ramp rates on organismal performance and fitness. Since we were not observing animals during this experiment, we used an opaque Styrofoam lid over the plate, but a clear plastic or glass lid could be used and would allow for experiments requiring continuous observation during a thermal ramp.

**Figure 3:**
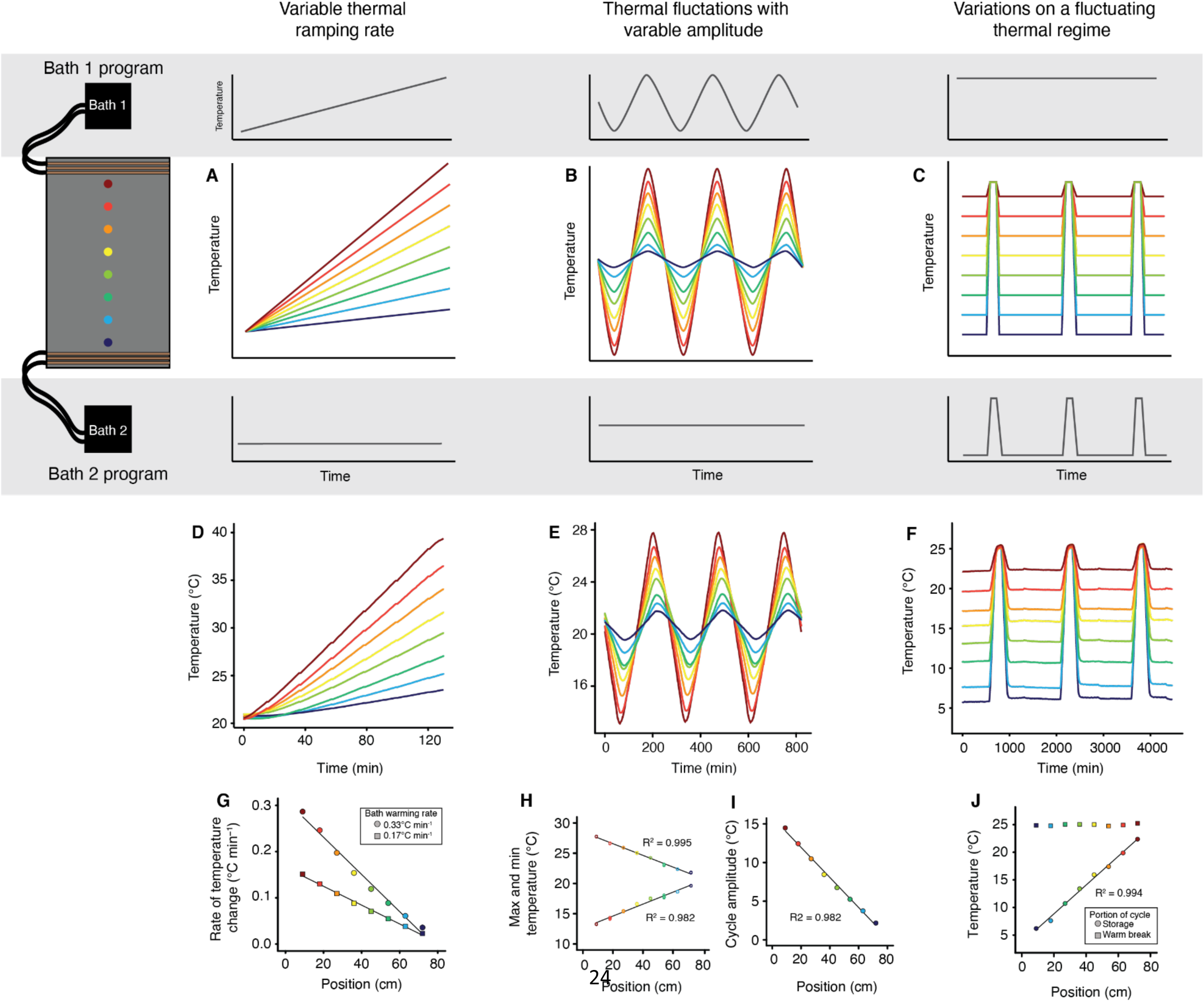
Dynamic thermal gradients allow for complex experiments using programmable refrigerated circulators. Programs shown produce predictable variations in warming rate, thermal cycle amplitude, or a chosen aspect of a fluctuating thermal regime over time. A-C) Expected relationships between time (x-axis) and temperature (y-axis) along the length of the gradient plate (color represents position across the length of the plate) when baths 1 and 2 are programmed as shown (grey background). D-F) Actual outcomes of bath programming experiments (temperature recordings over time). Each line represents the temperature of measurements at each position using type-K thermocouples. G) Rates of temperature change at each position during two runs using different warming rates of bath 2. This approach creates a linear relationship between position and thermal ramp rate. H-I) Maximum and minimum temperatures and cycle amplitude recorded using a cycling program. Cycling temperature of one bath (bath 1) while holding another constant (bath 2) produces repeatable cycle maxima and minima that vary by position across the plate and linearly alters cycle amplitude relative to the position. J) Relationship between temperature and position for cold storage temperature and warm break period in a fluctuating thermal regime simulation. A single aspect of the program can be altered predictably across the length of the plate (e.g., storage temp) while holding another aspect (warm break temperature) constant.

To create cycling temperatures, we set one circulator to cycle between high and low temperatures while the other was held at a constant temperature in the middle of cycle amplitude (Figure 3B). As expected, this set of programs produced predictable temperature cycles of varying amplitudes across the length of the plate (Figure 3E). The maxima and minima of each cycle changed linearly across the length of the plate (maxima R^2^ = 0.995, minima R^2^ = 0.982; Figure 3H), and thus the amplitude of the temperature variation varied linearly across the plate (R^2^= 0.982; Figure 3I). Thus, our system can be used to study the effects of thermal fluctuations on survival, growth, reproduction, or any other trait of interest. By testing the system setup while trialing sample placement, a carefully crafted range of variations in cycle amplitude can be created and used.

As a third and final test of the ability of our system to create dynamic thermal environments, we simulated a fluctuating thermal regime (FTR). FTR is characterized by a period of cool temperatures followed by a warm break and is often used to store insects for long periods for commercial purposes while slowing physiological aging and avoiding long term effects of low-temperature exposure (Figure 3C). To accomplish this, we held one bath at a constant temperature (+25°C) while the other bath completed cycles of 0°C with periodic warming to +25°C. As predicted (Figure 3C), this arrangement produced FTR cycles across the plate with a consistent temperature for the warm break (+25°C), while the cool “storage” temperatures varied linearly across the length of the plate (Figure 3F, J; R^2^=0.994). Thus, our design allows for careful manipulation of sample conditions following stepwise changes in the thermal program. In the case of FTR cycles, the sample storage or warm “break” temperature can be independently manipulated to study the effects of these treatments on insect survival or fitness. This specific application of our design would allow for high-throughput optimization of commercial or governmental insect storage programs.

### Proof of concept 1: Drosophila cold tolerance

We generated a low-temperature survival curve for *Drosophila* from a single use of the plate (Figure 4A; survival 2 h post-cold stress shown). We noted a sharp decrease in survival in males and females at approximately −2.2°C, and while the effect of sex was small (Fig. 4A), the large sample size contributed to both temperature (F_1,1578_=1945.8, P<0.001) and sex (F_1,1577_=84.6, P<0.001) having statistically significant effects on survival. This sex effect was driven by the fact that males were more tolerant of chilling at just one temperature point in the assay (−2.2°C; Figure 4A). At this temperature, nearly all males were alive 2 h after the cold stress while c. 70% of females were dead (Fig 4A). Survival was again measured at 24 h, and 48 h following removal from the cold exposure, so we tested whether survival outcomes changed between males and females in the 48 h following after exposure to −2.2°C (Figure 4B). Specifically, while males were far more likely to be scored as alive 2 h following the cold stress, they were far more likely to die in the ensuing 48 h (Figure 4B; mixed effects model recovery time x sex interaction: F_1,45_=7.34, *P*=0.002). While females never recovered from chill coma, males recovered the ability to stand, but then died. These results are similar to our recent report of latent chilling injury effects in virgin female flies following exposure to 0°C [39], which suggests that conditions leading to latent injury may be temperature-, sex-, and/or reproductive status-specific. Recording the same data with smaller sample size or fewer temperatures, over multiple rounds because of equipment limitations, or not recording survival over multiple time points could have missed these important differences between the sexes.

**Figure 4:**
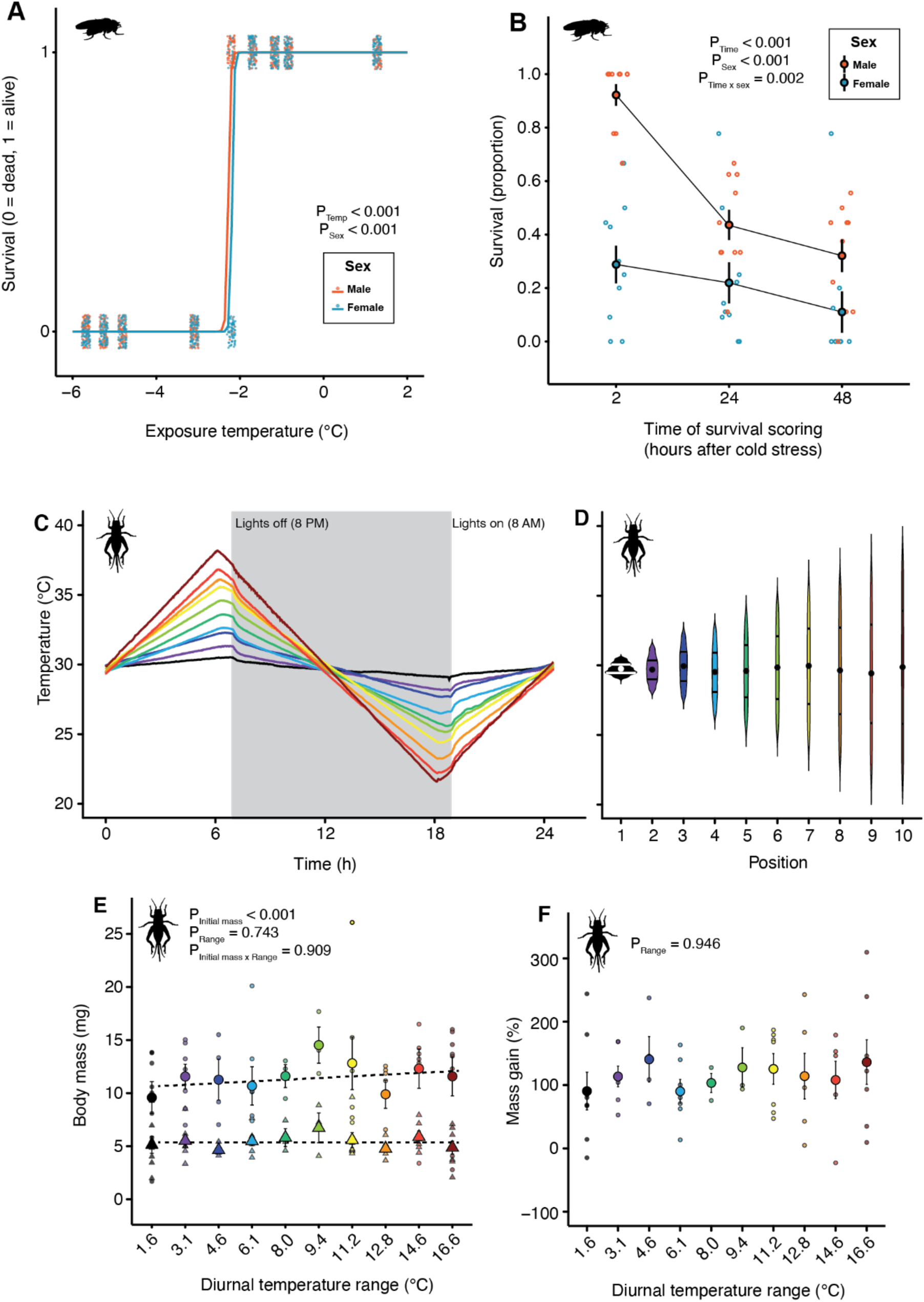
Proof of concept experiments to demonstrate the utility of the dynamic thermal gradient system. A) Survival of adult male (red) and female (blue) *D. melanogaster* following constant exposure to different temperatures for 4 h on the thermal plate. Each solid circle represents an individual fly. 1 = survived, 0 = dead. We noted that males were much more cold-tolerant when measured specifically at −2.21°C. B) Survival following exposure to −2.21°C for 4 h in the same flies across multiple rounds of survival assessment (2 h, 24 h, and 48 h). Although males appear more tolerant of chilling at first, they are more likely to suffer latent mortality. Each open circle represents an independent vial (containing ∼10 flies); solid circles represent the mean (± sem) survival. C) One day of the diurnal cycling program experienced by developing crickets (*Gryllodes sigillatus*). Different colours denote 10 different locations across the length of the plate with temperature ranges from 1.6°C (black) to 16.6°C (dark red). Small deviations in temperature can be seen when the lights switch on and off. D) Violin plots showing temperatures experienced by the crickets each day. Width denotes the amount of time spent at a given temperature. Solid circles denote the mean temperature experienced across the entire day (constant +30°C across the plate). Horizontal lines denote the 25% and 75% quartiles. E) Initial (triangles) and final (circles) mass of crickets at each location across the plate. Small symbols denote individual crickets and large symbols denote the mean ± sem. Variance in final body size was strongly related to initial size, but not significantly impacted by thermal variation. F) The same is true when adjusting for initial body size by expressing growth as a percentage of initial mass; thermal variability did not affect growth rates of *G. sigillatus*.

### Proof of concept 2: Gryllodes sigillatus growth during daily thermal cycles

To test the effects of thermal fluctuations on insect growth, we generated a 24-hour cycling regime from +40°C to +20°C (Figure 4C) that allowed us to create 10 different cycling regimes across the plate that varied in the amplitude of thermal fluctuation. We used this approach to record the growth rates of two-week old crickets over five days (Figure 4D). The starting weight of each cricket was measured before the start of the cycling regime and at the end (Figure 4E). We expected thermal variability to negatively impact growth rates. The initial mass of the cricket strongly predicted the final mass (F_1,116_=61.6, P<0.001), but surprisingly the position of the cricket on the plate (diurnal temperature range) did not affect final mass (F_9,107_ = 0.7, P = 0.743), or interact with initial mass to influence final mass (F_9,98_ = 0.4, P = 0.909). Most crickets approximately doubled in mass over the five-day period (Figure 4F). This implies that none of the cycling regimes created (ranging from 1.6 to 16.6°C of thermal range) resulted in any effect on growth rates.

### Conclusions

The thermal gradient system described and demonstrated here represents a powerful new approach to characterizing thermal performance in small organisms. This system can be built from readily available parts and with limited technical knowledge and can be used on a wide variety of sample types to measure nearly any thermal performance trait. The system produces stable static and/or predictable dynamic thermal gradients over a large surface area, permitting high throughput investigations. We are optimistic that this system as described or with creative improvements can enhance and accelerate research on the ecology, evolution, physiology, and molecular biology of ectothermic organisms, while also making these studies more economical for researchers with a limited budget.

## Supporting information

Data archive

## Acknowledgments

The authors wish to thank Charlie Reid and Hannah Davis for helpful discussion throughout this project, the staff at Entomo Farms for providing the crickets, and Matt Muzzatti for useful advice on caring for the crickets.

## Data Availability

All data described in this manuscript is provided as supplementary material.

